# On the automatic annotation of gene functions using observational data and phylogenetic trees

**DOI:** 10.1101/2020.05.14.095687

**Authors:** George G. Vega Yon, Duncan C. Thomas, John Morrison, Huaiyu Mi, Paul D. Thomas, Paul Marjoram

## Abstract

**Motivation:** Gene function annotation is important for a variety of downstream analyses of genetic data. Yet experimental characterization of function remains costly and slow, making computational prediction an important endeavor. In this paper we use a probabilistic evolutionary model built upon phylogenetic trees and experimental Gene Ontology functional annotations that allows automated prediction of function for unannotated genes.

**Results:** We have developed a computationally efficient model of evolution of gene annotations using phylogenies based on a Bayesian framework using Markov Chain Monte Carlo for parameter estimation. Unlike previous approaches, our method is able to estimate parameters over many different phylogenetic trees and functions. The resulting parameters agree with biological intuition, such as the increased probability of function change following gene duplication. The method performs well on leave-one-out validation, and we further validated some of the predictions in the experimental scientific literature.

**Availability:** Our method has been implemented as an R package and it is available online at https://github.com/USCBiostats/aphylo. Code needed to reproduce the tables and figures can be found in https://github.com/USCbiostats/aphylo-simulations.

**Author summary:** Understanding the individual role that genes play in life is a key issue in biomedical-sciences. While information regarding gene functions is continuously growing, the number of genes with unknown biological purpose is yet greater. Because of this, scientists have dedicated much of their time to build and design tools that automatically infer gene functions. In this paper, we present yet another attempt to do such. While very simple, our model of gene-function evolution has some key features that have the potential to generate an impact in the field: (a) compared to other methods, ours is highly-scalable, which means that it is possible to simultaneously analyze hundreds of what are known as gene-families, compromising thousands of genes, (b) supports our biological intuition as our model’s data-driven results coherently agree with what theory dictates regarding how gene-functions evolved, (c) notwithstanding its simplicity, the model’s prediction accuracy is comparable to other more complex alternatives, and (d) perhaps most importantly, it can be used to both support new annotations and to suggest areas in which existing annotations show inconsistencies that may indicate errors or controversies.

## 1 Introduction

The overwhelming majority of sequences in public databases remain experimentally uncharacterized, a trend that is increasing rapidly with the ease of modern sequencing technologies. To give a rough idea of the disparity between characterized and uncharacterized sequences, there are ~ 15 million protein sequences in the UniProt database that are candidates for annotation, while, only 81,000 (0.3%) have been annotated with a Gene Ontology (GO) term based on experimental evidence. It is therefore a high priority to develop powerful and reliable computational methods for inferring protein function. Many methods have been developed, and a growing number of these have been assessed in the two Critical Assessment of Function Prediction (CAFA) experiments held to date [27, 19].

In previous work, we developed a semi-automated method for inferring gene function based on creating an explicit model of function evolution through a gene tree [13]. This approach adopts the “phylogenetic” formulation of function prediction first proposed by Eisen [6], and the use of GO terms to describe function as implemented in the SIFTER software (Statistical Inference of Function Through Evolutionary Relationships) developed by Engelhardt et al. [8]. To date, our semi-automated method has been applied to over 5000 distinct gene families, resulting in millions of annotations for protein coding genes from 142 different fully sequenced genomes. However, this approach requires manual review of GO annotations, and manual construction of distinct models of gene function evolution for each of the 5000 families. Even using extensive curation and complex software, called PAINT, the semi-automated inference process has taken multiple person-years. Further, the semi-automated process cannot keep up with the revisions that are constantly necessary due to continued growth in experimentally supported GO annotations.

Here, we describe an attempt to develop a fully automated, probabilistic model of function evolution, that leverages the manually constructed evolutionary models from PAINT, for both training and assessing our new method. Related work has previously been undertaken in this area, and a probabilistic framework for function prediction has been implemented in SIFTER. However, this framework assumes a model of function evolution that limits its applicability in function prediction. First, it was developed specifically only to treat molecular function, and not cellular component and biological process terms that could, in principle, be predicted in a phylogenetic framework. Cellular component and biological process GO terms are the most commonly used in most applications of GO [29], so these are of particular practical importance. And second, while it has proven to provide good predictions (about 73% accuracy using the area under the curve statistic), it cannot be scaled and the model itself provides no theoretical insights whatsoever. In practice, perhaps the most serious problem for any inference method is the sparseness of experimental annotations compared to the size of the tree. As a result, standard parameter estimation techniques for the SIFTER framework are impossible, and consequently for SIFTER2.0, the transition matrix parameters are fixed at somewhat arbitrary values [7].

In order to overcome these problems, we propose a much simpler evolutionary model, in which (like the semi-automated PAINT approach) each function is treated as an independent character that can take the value 1 (present) or 0 (absent) at any given node in the phylogenetic tree. Using information about experimental annotations available in the GO database, and phylogenetic trees from the PANTHER project [22], we show that we can build a parsimonious and highly scalable model of functional evolution that provides both intuitive insights on the evolution of functions and highly accurate predictions.

The content of the paper is as follows: section 2 presents mathematical notation and formally introduces the model, including parameter estimation and calculation of posterior probabilities. Section 3 presents a large-scale application in which we take a sample of annotations from the GO database along with their corresponding phylogenies, fit our model, and analyze the results. Finally, section 4 discusses method limitations and future research directions for this project.

## 2 Methods

In general terms, we propose a probabilistic model that reflects the process by which gene functions are propagated through an evolutionary tree. The fundamental idea is that for any given node in the tree, we can write down the probability of observing a function to be present for the gene as a function of model parameters and the functional state of its parent node, essentially modeling probability of gaining and losing function.

The baseline model has 5 parameters: the probability that the most recent common ancestor of the entire tree, i.e. the root node, had the function of interest, the probability of functional gain (the offspring node gains the function when the parent node did not have it), the probability of functional loss (the offspring node loses the function when the parent node had it), and two additional parameters capturing the probability that the gene was incorrectly labeled, i.e. the probability of classifying an absent function as present and vice-versa. We also consider two simple extensions: specifying the functional gain and loss by type of evolutionary event (speciation or duplication), and pooling information across trees.

As explained later, in this version of our model, if there are multiple gene functions of interest, we analyze one function at a time (i.e., we treat those functions as independent). But later we also show results for a joint analysis of multiple functions, assuming they share parameter values. We then discuss further extensions to our model in the Discussion section.

We assume that our starting data consists of a set of gene annotations for a given tree. We further assume that those annotations occur only at the leaves of the tree (as is typical) and that those leaves were annotated via manual curation derived from experimental data (and are therefore subject to the misclassification probabilities outlined above). Our goal is then to predict the functional state of un-annotated leaves (and, conceivably, to assess the likely quality of the existing annotations). Our perspective is Bayesian. Therefore, we proceed by estimating the posterior distributions of the parameters of the evolutionary model and then, conditional on those estimated distributions, computing the posterior probability of function for each node on the tree.

We now give full details, beginning with some notation.

### 2.1 Definitions – Annotated Phylogenetic Tree

A phylogenetic tree 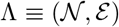 is a tuple of nodes 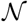, and edges 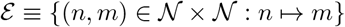 defined by the binary operator ↦ *parent of*. We define 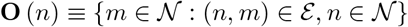 as the set of offspring of node *n*, and 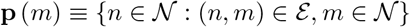 as the parent node of node *m*. Given a tree Λ, the set of leaf nodes is defined as 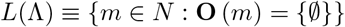.

Let 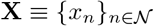 be a vector of annotations in which the element *x_n_* denotes the state of the function at node n, taking the value 1 if such function is present, and 0 otherwise. We define an Annotated Phylogenetic Tree as the tuple *D* ≡ (Λ, **X**).

Our goal is to infer the true state of **X**, while only observing an imperfect approximation of it, 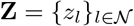, derived from experimentally supported GO annotations [29]. Typically only a small subset of the leaf nodes will have been annotated. We refer to the tuple 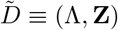 as an Experimentally Annotated Phylogenetic Tree. Finally, let 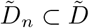 be the induced experimentally annotated subtree that includes all information – i.e., tree structure and observed annotations – regarding the descendants of node *n* (including node *n* itself). This object constitutes a key element of the likelihood calculation. Table 1 summarizes the mathematical notation.

**Table 1.**
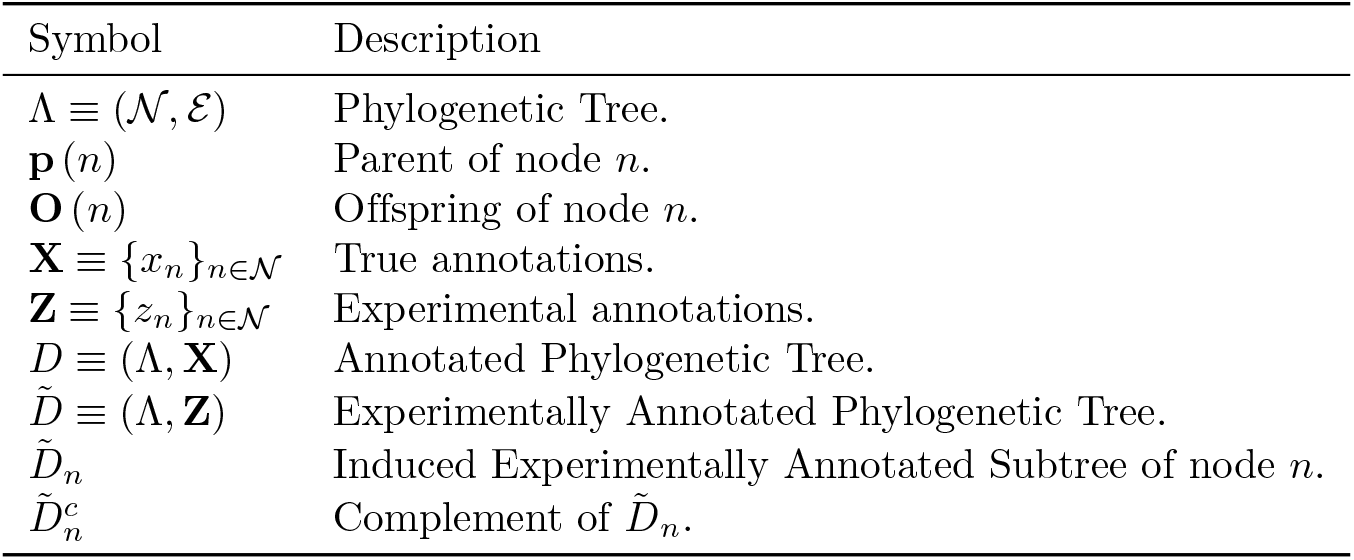
Mathematical Notation

### 2.2 Likelihood of an Annotated Phylogenetic Tree

#### 2.2.1 Baseline model

Our evolutionary model for gene function is characterized by the phylogeny and five model parameters: the probability that the root node has the function, *π*, the probability of an offspring node either gaining or losing a function, *μ*_01_ and *μ*_10_, and the probability that either an absent function is misclassified as present or that a present function is misclassified as absent, *ψ*_01_ and *ψ*_10_ respectively, in the input data.

To simplify notation, we will write *ψ* = (*ψ*_01_, *Ψ*_10_) and *μ* = (*μ*_01_,*μ*_10_) when referring to those pairs. Table 2 summarizes the model parameters.

**Table 2.** Model parameters

| Parameter | Description |
| --- | --- |
| $\pi$ | Probability that the root node has the function |
| $\mu_{01}$ | Probability of gaining a function |
| $\mu_{10}$ | Probability of losing a function |
| $\psi_{01}$ | Probability of experimental mislabeling of a 0 |
| $\psi_{10}$ | Probability of experimental mislabeling of a 1 |

Following [10], we use a pruning algorithm to compute the likelihood of an annotated phylogenetic tree. In doing so, we visit each node in the tree following a post-order traversal search, i.e., we start at the leaf nodes and follow their ancestors up the tree until we reach the root node.

Given the true state of leaf node *l*, the mislabeling probabilities are defined as follows:

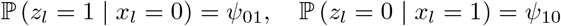

We can now calculate the probability of leaf *l* having state *z_l_*, given that its true state is *x_l_* as:

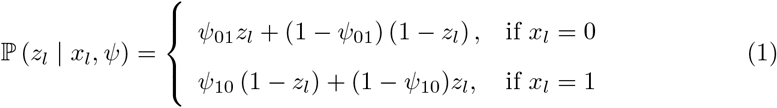

Similarly, the functional gain and functional loss probabilities are defined as follows:

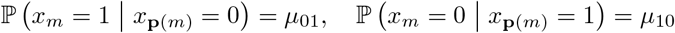

Note that in this version of our model we assume that these probabilities are independent of the time that has passed along the branch connecting the two nodes. We return to this point in the Discussion. Now, for any internal node 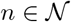, we can calculate the probability of it having state *x_n_*, given the state of its parent **p** (*n*) and the vector of parameters *μ*, as:

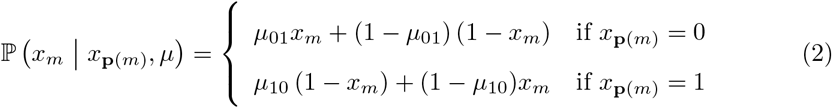

Together with (1), and following [10], this allows us to calculate the probability of the interior node *n* having state *x_n_* conditional on 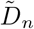, its induced subtree, as the product of the conditional probabilities of the induced subtrees of its offspring:

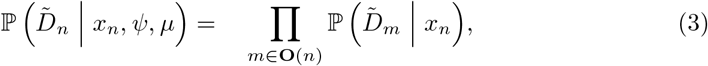

where

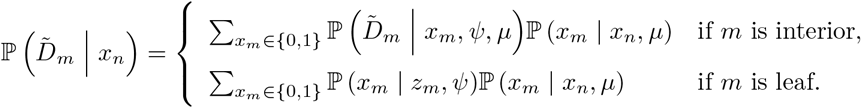

This is a recursive function that depends upon knowing the offspring state probabilities for internal nodes, which, since we are using a pruning algorithm, will already have been calculated as part of the process. Finally, the probability of the experimentally annotated phylogenetic tree can be computed using the root node conditional state probabilities:

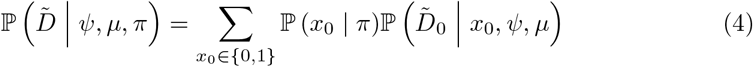

where 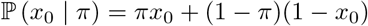.

In the next section we introduce additional refinements that allow us take into account prior biological knowledge that might constrain the parameter space and alleviate the typical sparseness of the curated data – we return to these issues in the Discussion.

#### 2.2.2 Pooled data models

As mentioned earlier, the sparsity of experimental annotations typically observed in this context makes inference challenging. However, in order to improve inference we might attempt to “borrow strength” across a set of functions, by combining them in a single analysis. As a “proof of principle” of this idea, we will also show results in which we assume that sets of annotated phylogenetic trees share population parameters. This way, we can estimate a joint model by pooling those annotated phylogenetic trees, providing stronger inference about the evolutionary parameters and therefore, we hope, more accurate inference of gene annotations. We note that we have strived to make sure our software implementation extremely computationally efficient, allowing us to estimate a joint model with hundreds of annotated trees in reasonable time on a single machine.

Our results show that using a pooled-data model for parameter estimation greatly increases the model’s predictive power. In the Discussion we will outline possible future, less simplified, approaches to this problem.

#### 2.2.3 Type of evolutionary event

The non-leaf nodes in the phylogenetic trees that we are considering here come in two types, reflecting duplication and speciation events. It has been widely-observed that change of gene function most commonly occurs after duplication event (broadly speaking, the extra copy of a gene is free to change function because the original copy is still present.) Speciation events, on the other hand, are mostly driven by external causes that do not necessarily relate to functional changes in one specific gene, meaning that, while we may observe some differences between siblings, their function generally remains the same.

Our model reflects this biological insight by allowing the functional gain and loss probabilities to differ by type of event. In particular, instead of having a single parameter pair (*μ*_01_, *μ*_10_), we now have two pairs of functional gain and loss parameters, (*μ*_01*d*_, *μ*_10*d*_) and (*μ*_01*s*_, *μ*_10*s*_), the gain and loss probabilities for duplication and speciation events, respectively.

### 2.3 Estimation and Prediction

We adopt a Bayesian framework, introducing priors for the model parameters. In particular, as described in section 3, we use Beta priors for all model parameters. Estimation is then performed using Markov chain Monte Carlo [MCMC] using an Adaptive Metropolis transition kernel [14] with reflecting boundaries at [0,1] (see, for example, [32]) as implemented in the *fmcmc* R package [30]. We do this using the R package *aphylo* which we have developed to implement this model (https://github.com/USCbiostats/aphylo). The R package also allows estimating model parameters using Maximum Likelihood Estimation [MLE] and Maximum A Posteriori estimates [MAP].

Regarding model predictions, once we fit the model parameters we use the calculated probabilities during the computation of the likelihood function (pruning process) and feed them into the posterior probability prediction algorithm, which is exhaustively described in subsection 5.1 (see also [18]). It is important to notice that, in general, predictions are made using a leave-one-out approach, meaning that to compute the probability that a given gene has a given function, we remove the annotation for that gene from the data before calculating the overall likelihood of the tree. Otherwise we would be including that gene’s own observed annotation when predicting itself. The latter point is relevant for evaluating prediction accuracy. Prediction of unannotated genes, on the other hand, is performed using all the available information.

In general, whenever using MCMC to estimate the model parameters, we used 4 parallel chains and verified convergence using the Gelman-Rubin [3] statistic. In all cases, we sampled every tenth iteration after applying a burn-in of half of the sample. As a rule of thumb, the processes were considered to have reached a stationary state once the potential scale reduction factor statistic was below 1.1. When we report point estimates for parameters we use the mean value across sampled iterations for all chains after burn-in.

To measure prediction quality, we use the Mean Absolute Error [MAE], which we calculate as follows:

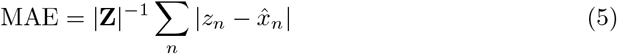

Where |*Z*| is the number of observed annotations, *z_n_* is the observed annotation (0/1), and 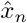 is the predicted probability that the *n*-th observation has the function of interest. Since *z_n_* is binary, perfect predictions result in a score of 0, completely wrong predictions result in a score of 1, and a random coin toss results in a score of 0.5.

While other statistics, such as accuracy and Area Under the Receiver Operating Characteristic Curve [AUCR] (see [9] for a general overview), are most commonly used for binary prediction models, we prefer to use MAE since it is: (a) robust to unbalanced data, (b) probabilistic and thus does not depend on choice of threshold for prediction, and (c) robust to probabilistic noise, see [11, 5] for a more general discussion.

We now turn our attention to an application using experimental annotations from the Gene Ontology Project.

## 3 Results

To evaluate performance of our model, we used data from the Gene Ontology project. In particular, in order to assess the potential utility of combining information across similar genes, we used our model in two different ways: a pooled-data model, in which we treated all trees as if they had the same evolutionary parameter values, and a one-tree-at-a-time model, in which we estimated parameters, and then imputed functional status, for each tree separately.

To reduce sparsity of annotations somewhat, from the entire set of available annotated phylogenies in PANTHER [22] (about 15,000), we selected only those in which there were at least two annotations of each type (i.e., at least two genes annotated as possessing a given function, and two as lacking the function). This yielded a total of 141 experimentally annotated trees. A key feature of the resulting dataset is that most leaves still have no annotation, as experimental evidence is still very sparse. Furthermore, in order to test the robustness of our results, we compared 4 analyses: MCMC with a uniform prior for parameter values, MCMC with a Beta prior (for both of which we report the mean,) Maximum Likelihood Estimation [MLE], and Maximum A Posteriori estimates [MAP]. The shape parameters used for MCMC and MAP with Beta priors, which are detailed in Table 3, were chosen to reflect the biological intuition that change of state probabilities should be higher at duplication events.

**Table 3.** Parameters for the Beta priors used in MCMC and MAP estimation. Shape parameters (*α, β*) were set with the prior that functional gain and loss, (*μ*_01*d*_, *μ*_10*d*_), are more likely at duplication events.

| Parameter | $\alpha$ | $\beta$ | Mean<br>$\alpha/(\alpha + \beta)$ |
| --- | --- | --- | --- |
| $\psi_{01}, \psi_{10}$ | 2 | 9 | 0.18 |
| $\mu_{01d}, \mu_{10d}$ | 9 | 2 | 0.81 |
| $\mu_{01s}, \mu_{10s}$ | 2 | 9 | 0.18 |
| $\pi$ | 2 | 9 | 0.18 |

Finally, accuracy was measured using leave-one-out cross-validated Mean Absolute Error [MAE].

### 3.1 Pooled-data model

First we show results for the pooled-data model, in which we combine the analysis of multiple trees, assuming parameter values do not vary between trees. The resulting parameter estimates are presented in Table 4, and are seen to reflect biological intuition: we see high probabilities of change of function at duplication events, which occurs regardless of whether we use an informative prior or not. While both probabilities are high, gain of function is roughly twice as likely as loss of function at such nodes. We note that we estimate relatively high values of *ψ*_01_ and low values for *ψ*_10_. We believe this is largely due to the sparsity of 0, i.e. *“absent*”, experimental annotations, which means that high values for *ψ*_10_ are implausible and implies that any mislabelings must perforce be that of assigning function falsely. Thus we think the relatively high estimates of *ψ*_01_ are an artefact that will likely disappear in the presence of less sparse annotation data.

**Table 4.**
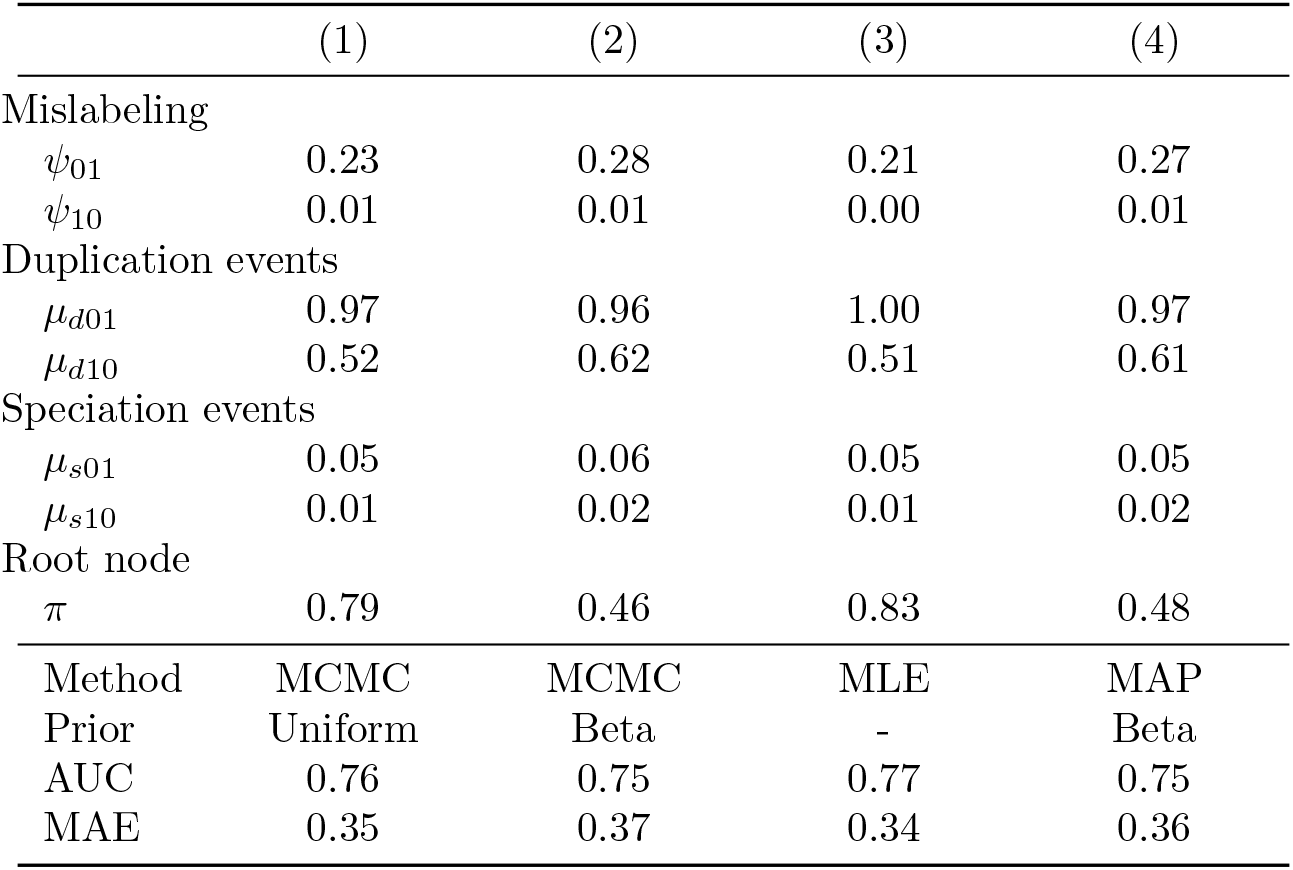
Parameter estimates for model with experimentally annotated trees. Overall, the parameter estimates show a high level of correlation across methods. Regardless of the method, with the exception of the root node probabilities, most parameter estimates remain the same. As expected, mutation rates at duplication nodes, indexed with a *d*, are higher than those observed in speciation nodes, indexed with an *s*.

We note that parameter estimates are consistent across estimation methods. Here we used both non-informative (uniform) and relatively informative priors. We see that choice of prior had no significant effect on the final set of estimates, meaning that these are driven by data and not by the specified priors. The only exception is for the root node, *π*, which is hard to estimate since the root node exists far back in evolutionary time [12]. We also conducted a simulation study across a broader range of parameter values, using simulated datasets, in which we saw similar behavior. These results are shown in subsection 5.2. However, we note that when analyzing one tree at a time, this robustness is lost [results not shown] because the sparsity of data typical in any single given tree does not permit strong inference regarding evolutionary parameters. Therefore, the prior becomes more influential.

We note that each instance of the estimation process (reaching stationary distribution in MCMC or finding the maxima in MLE and MAP) took between 1 to 3 minutes using a regular laptop computer. To put this number into context, we use the time calculator provided by SIFTER [8, 7], which is available at http://sifter.berkeley.edu/complexity/, as a benchmark. Since, as of today, SIFTER is designed to be used with one tree at a time, we compared our run time with the estimated run time for SIFTER for a tree with 590 leaves (the average size in our data). In such case, they estimate their algorithm would take about 1 minute to complete, with an upper bound of ~9 minutes, which translates into about 141 minutes to run on our entire dataset (upper bound of ~21 hours). However, in SIFTER’s defense, we note that their model allows for a greater level of generality than does our current implementation of our own method. Specifically, they allow for correlated changes across multiple gene functions. In the simplest case, using “truncation level” 1 as defined in their paper, the estimated run time for an analysis comparable in size with the one conducted here, 141 functions and 83,000 leaves, is 6 hours. However, for truncation level 2, the estimated run time is 2.2 years. In the Discussion section we outline how we propose to extend our method in future to allow for for more sophisticated joint analyses across function than the one we present here.

### 3.2 One-tree-at-a-time

The joint analysis of trees we describe in the previous section makes the enormous assumption that parameter values do not vary across trees. This assumption is made for pragmatic reasons: while it is not likely to be correct (and more general approaches will be described in the Discussion), it is a simple analysis to implement and it allows us to quickly assess a baseline for the improvement in overall prediction accuracy that might result from performing inference jointly across trees. So, in this section we report results of a one-at-a-time analysis of gene function and assess whether the pooled model outperforms a one-at-a-time approach with regards to accuracy, even using our extremely restrictive assumptions.

For each phylogeny, we fitted two different models, one with an uninformative uniform prior and another with the same beta priors used for the pooled-data models, this is, beta priors with expected values close to zero or one, depending on the parameter. Again, as before, model predictions were undertaken in a leave-one-out approach and corresponding MAEs were calculated for each set of predictions. Table 5 shows the differences in Mean Absolute Error [MAE] between the various approaches.

**Table 5.** Differences in Mean Absolute Error [MAE]. Each cell shows the 95% confidence interval for the difference in MAE resulting from two methods (row method minus column method). Cells are color coded blue when the method on that row has a significantly small MAE than the method on that column; Conversely, cells are colored red when the method in that column outperforms the method in that row. Overall, predictions calculated using the parameter estimates from *pooled-data* predictions outperform *one-at-a-time.*

|  | *Pooled-data* | One-at-a-time |  |
| --- | --- | --- | --- |
|  | Beta prior | Unif. prior | Beta Prior |
| *Pooled-data* |  |  |  |
| Unif. prior | [-0.02,-0.01] | [-0.14,-0.10] | [-0.06,-0.03] |
| Beta prior | - | [-0.12,-0.09] | [-0.04,-0.01] |
| *One-at-a-time* |  |  |  |
| Unif. prior | - | - | [ 0.06, 0.09] |

From Table 5 we can see three main points: First, in the case of the one-at-a-time predictions, informative priors have a significantly positive impact on accuracy.

Compared to the predictions using beta priors, predictions based on the MAE estimates with uniform priors have a significantly higher MAE with a 95% confidence interval [c.i.] ranging [0.06, 0.09]. Second, predictions based on the pooled-data estimates outperform predictions based on one-at-a-time parameter estimates, regardless of the prior used. And third, in the case of predictions based on pooled-data estimates, model predictions based on the parameter estimates using a uniform prior have significantly smaller MAE than predictions based on parameter estimates using a beta prior with a 95% c.i. equal to [−0.02, −0.01].

Given the sparsity of experimental annotation on most trees, a natural question to ask is: How important is it to have good quality parameter estimates in order to achieve good-quality predictions of gene function? In the next section we perform a sensitivity analysis to address this question.

### 3.3 Sensitivity Analysis

A fundamental question to ask is how much the accuracy of the posterior probabilities for gene function depends on the inferred values of the underlying evolutionary parameters used to make the predictions. Figure 1 shows how the distribution of the Mean Absolute Error [MAE] changes as a function of using parameters different from those obtained in the fitting process. In particular, each block of boxplots shows a *ceteris paribus* type of analysis, this is, MAE as a function of changing one parameter at a time while fixing the others.

**Fig 1.**
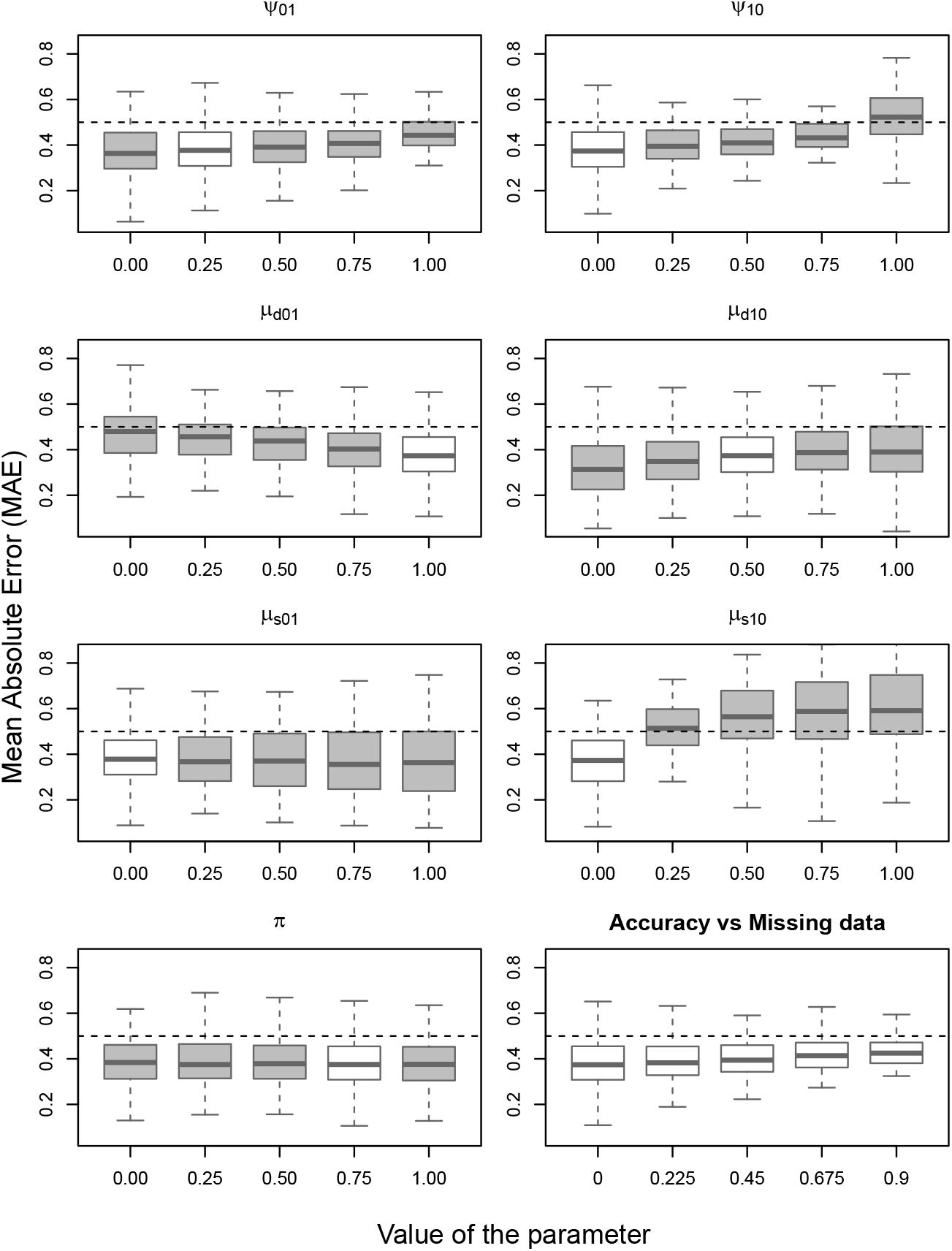
Sensitivity analysis. Boxplots of MAEs as a function of single parameter updates. The first seven plots show how the MAEs change as a function of fixing the given parameter value raging from 0 to 1. In each of these plots, the white box indicates the parameter value used to generate the data (i.e., the ‘‘correct” parameter value). The last boxplot shows the distribution of the MAEs as a function of the amount of available annotation data, that is, how accuracy changes as the degree of missing annotations increases.

In the case of the mislabeling probabilities (first row), it is not until they reach values close to one that the prediction accuracy significantly deteriorates. While higher values of *ψ*_10_ have a consistent negative impact on accuracy, the same is not evident for *ψ*_01_. The latter could be a consequence of the low prevalence of 0 (“no function”) annotations in the data, as illustrated in Figure 2.

**Fig 2.**
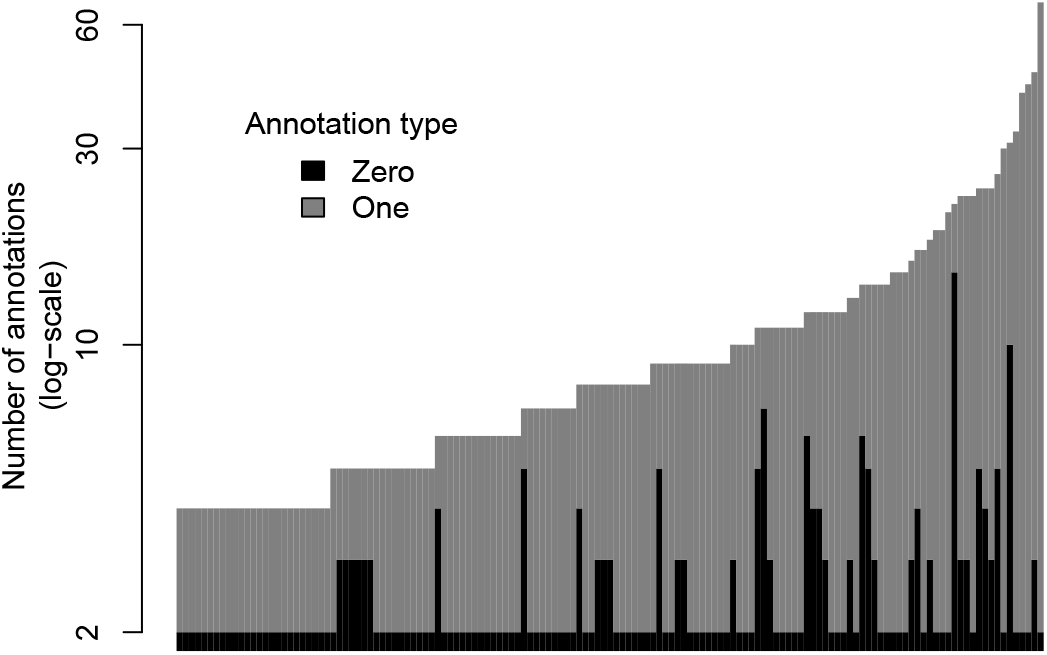
Number of annotated leaves for all 141 trees by type of annotation. Most of the annotations available in GO are positive assertions of function, that is, 1s. Furthermore, as observed in the figure, all of the trees used in this study have two “Not” (i.e., 0) annotations, which is a direct consequence of the selection criteria used, as we explicitly set a minimum of two annotations of each type per tree+function.

Functional gain and loss probabilities (second and third rows) have a larger effect on MAEs when misspecified, especially at speciation events, (*μ*_*s*01_, *μ*_*s*10_). This is largely because speciation events are much more common than duplication events across trees. Figure 3 shows a detailed picture of the number of internal nodes by type of event. Of the seven parameters in the model, the root node probability π has a negligible impact on MAE.

**Fig 3.**
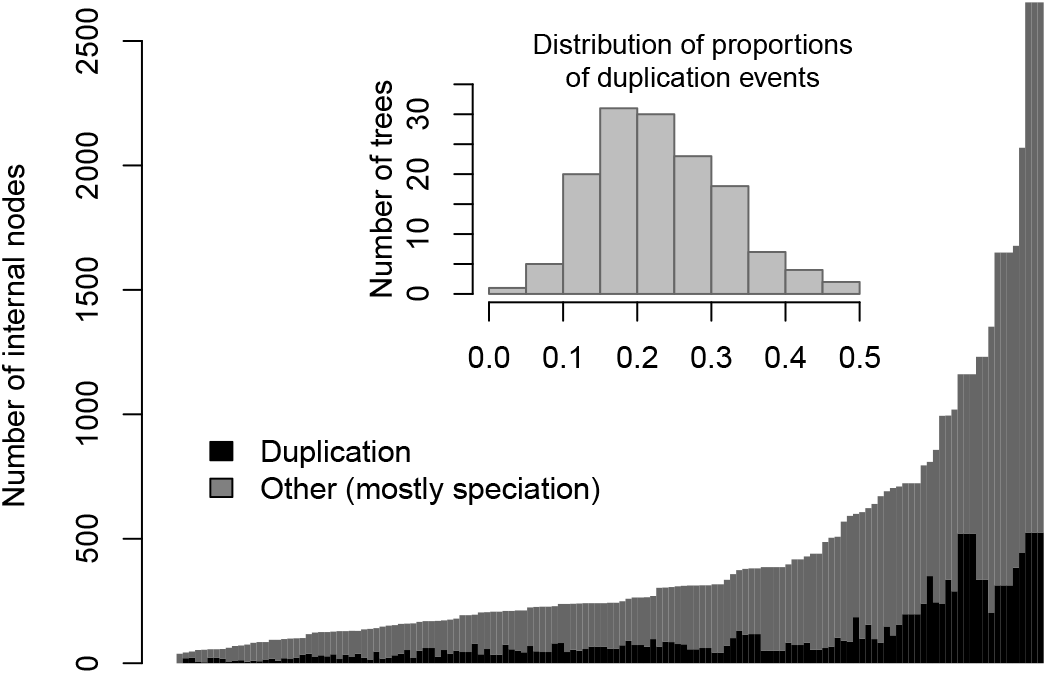
Number of internal nodes by type of event. Each bar represents one of the 141 trees used in the paper by type of event (mostly speciation). The embedded histogram shows the distribution of the prevalance of duplication events per tree. The minority of the events, about 20%, are duplications.

Finally, we evaluated the impact of removing annotations on accuracy. While a significant drop may not be obvious from the last box in Figure 1, a paired t-test found a significant drop in the accuracy levels.

### 3.4 Featured examples

An important determinant of the quality of the predictions is how the annotations are co-located. For the model to be able to accurately predict a 1, it is necessary either that it follows immediately after a duplication event in a neighborhood of zeros (so that function can gained with reasonable probability), or that a group of relatives within a clade have the function (i.e., its siblings have the function). Figures 4 and 5 show examples of high and low accuracy predictions respectively, illustrating how co-location impacts accuracy.

**Fig 4.**
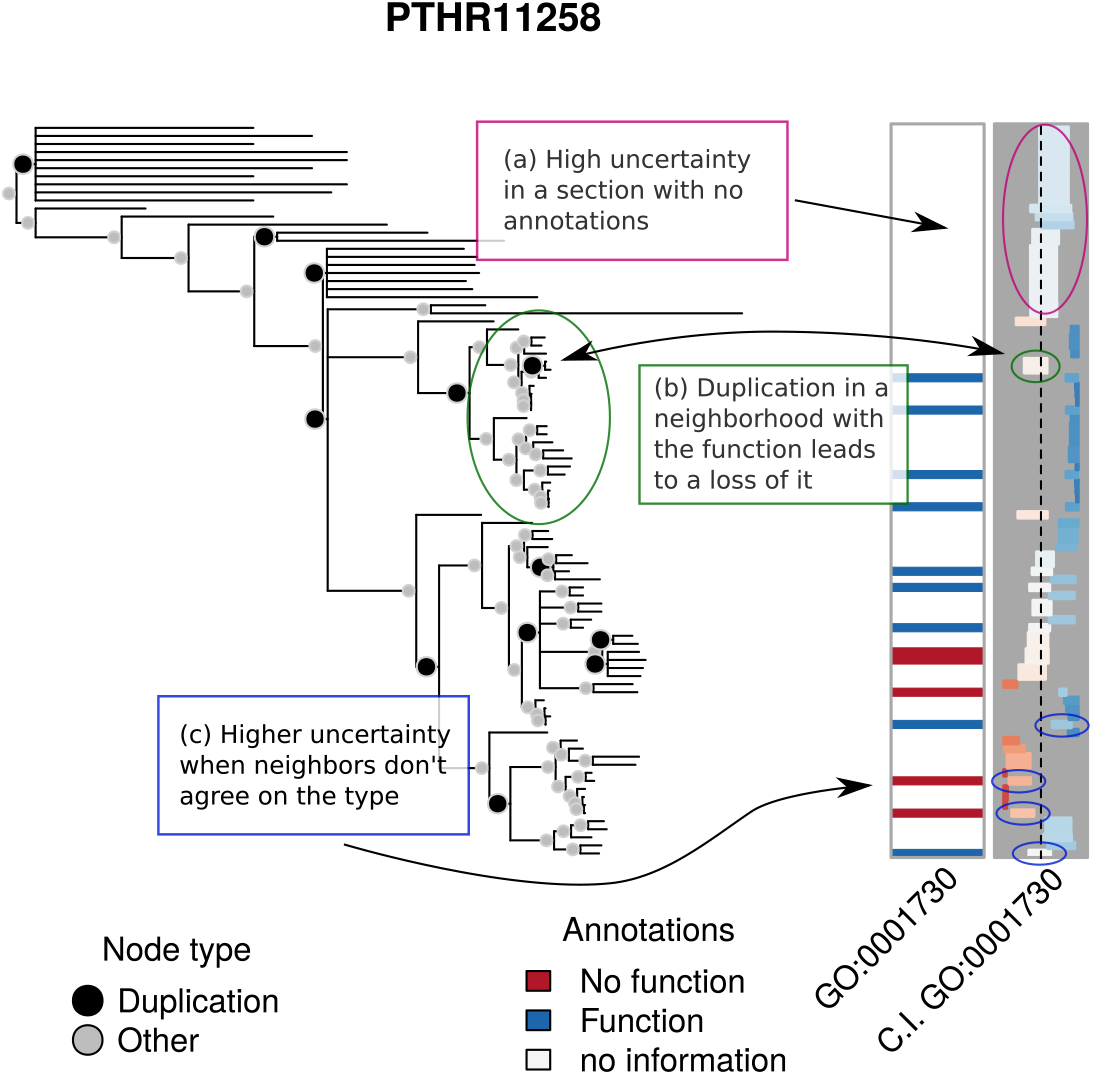
High accuracy predictions. The first set of annotations, first column, shows the experimental annotations of the term GO:0001730 for PTHR11258. The second column shows the 95% C.I. of the predicted annotations. The column ranges from 0 (left end) to 1 (right-end). Bars closer to the left are colored red to indicate that lack of function is suggested, while bars closer to the right are colored blue to indicate function is suggested. Depth of color corresponds to strength of inference. The AUC for this analysis is 0.91 and the Mean Absolute Error is 0.34.

**Fig 5.**
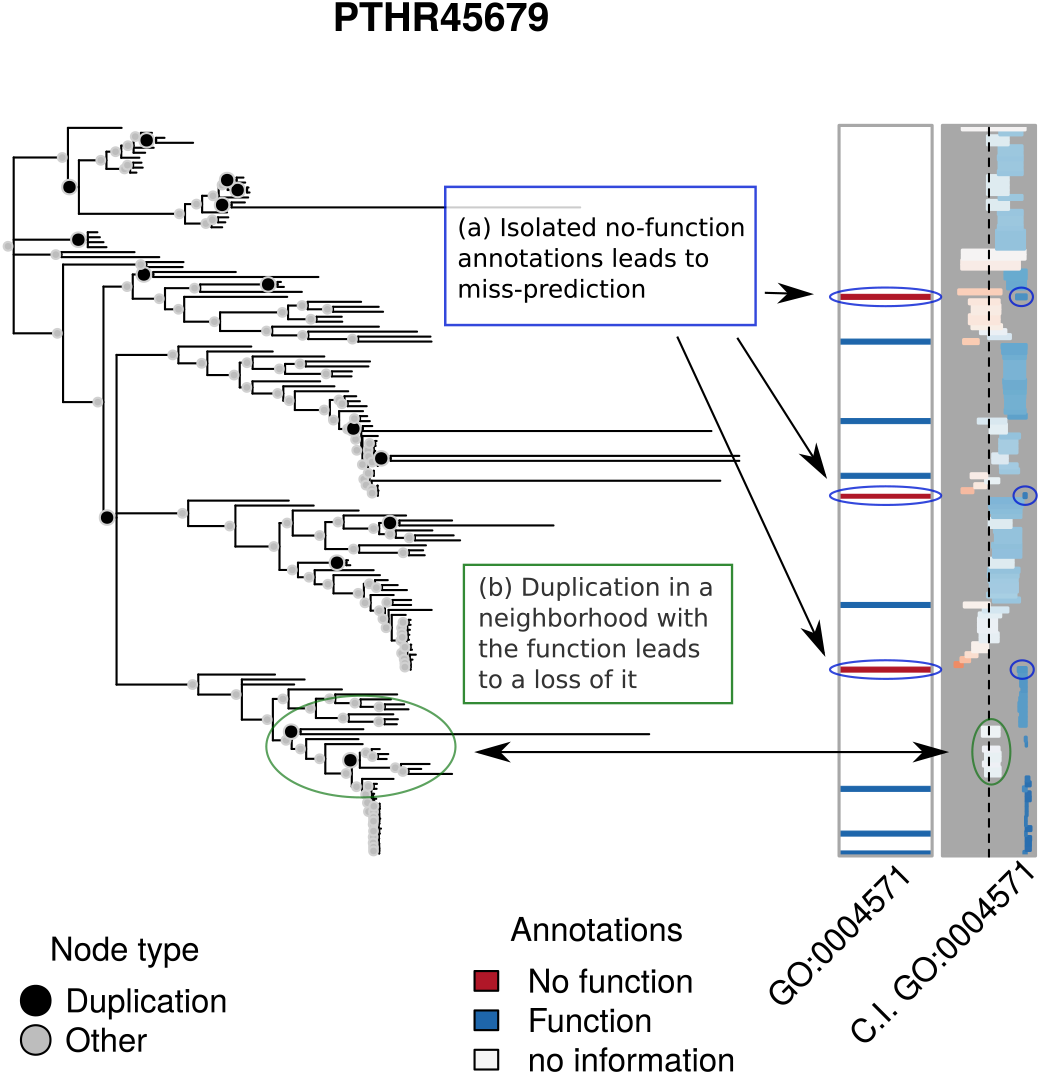
Low accuracy predictions. The first set of annotations, first column, shows the experimental annotations of the term GO:0004571 for PTHR45679. The second column shows the 95% C.I. of the predicted annotations using leave-one-out. Bars closer to the left are colored red to indicate that lack of function is suggested, while bars closer to the right are colored blue to indicate function is suggested. Depth of color corresponds to strength of inference. The AUC for this analysis is 0.33 and the Mean Absolute Error is 0.52.

In the previous analyses, we reported predictions based on point estimates. It is more informative to leverage the fact that we have posterior distributions for parameter values. Hence, in this section, in which we are looking in detail at specific trees, we exploit the full posterior distribution, and generate credible intervals by repeatedly sampling parameter values from their posterior distribution and then imputing function using each set of sampled parameter values. As before, we used a leave-one-out approach to make predictions, meaning that to predict any given leaf, we removed any annotation for that leaf and recalculated the probabilities described in subsection 5.1.

In the high accuracy example, Figure 4, which predicts the go term GO:0001730 on PANTHER family PTHR11258, we highlight three features of the figure. First, in box (a), the confidence intervals for the posterior probabilities in sections of the tree that have no annotations tend to be wider, which is a consequence of the degree to which posterior probabilities depend upon having nearby annotated nodes. Second, duplication events serve as turning points for function, as they can either trigger a functional loss, as highlighted in box (b), or a functional gain. The available data within the clade sets the overall trend for whether a function is present or not. The clade in box (b) is particularly interesting since there is only one duplication event and all of the annotated leaves have the function, which triggers a potential loss for the descendants of the duplication event. Finally, in regions in which labels are heterogeneous, it is harder to make a decisive prediction, as should be the case. This is why, in the leave-one-out predictions, the highlighted predictions in box (c) show wider confidence intervals and posterior probabilities closer to 0.5 than to the true state.

Figure 5 illustrates one way in which low accuracy can result. In particular, as highlighted by box (a), while the number of annotations is relatively large, (more than 10 over the entire tree, see Figure 2), the ‘absent’ (zero) annotations are distributed sparsely across the tree, having no immediate neighbor with an annotation of the same type. In fact, in every case, their closest annotated neighbors have the function. As a result, all zero-type annotations are wrongly labelled, with the model assigning a high probability of having the function. Similar to what is seen in the high accuracy case, duplication events within a clade with genes of mostly one type have a large impact on the posterior probabilities, as indicated by box (b).

In an effort to understand why the predictions were so poor in this case, we reviewed the family in detail. Because all GO annotations include references to the scientific publications upon which they are based, we were able to review the literature for genes in this family, the ER degradation enhancing mannosidase (EDEM) family. It turns out that the inconsistency of the annotations in the tree reflect a bona fide scientific controversy [24], with some publications reporting evidence that these proteins function as enzymes that cleave mannose linkages in glycoproteins, while others report that they bind but do not cleave these linkages. This explains the divergent experimental annotations among members of this family, with some genes being annotated as having mannosidase activity (e.g. MNL1 in budding yeast), others as lacking this activity (e.g. mnl1 in fission yeast), and some with conflicting annotations (e.g. human EDEM1). The inability of our model to correctly predict the experimental annotations actually has a practical utility in this case, by identifying an area of inconsistency in the GO knowledgebase.

### 3.5 Discoveries

We now explore to what extent predictions from our model can be used to suggest function. Using the estimates from the pooled-data model with uniform priors, we calculated posterior probabilities for all the genes involved in the 141 GO trees+functions analyzed. Moreover, we calculated 95% credible intervals for each prediction using the posterior distribution of the parameters and from there obtained a set of potential new annotations.

While all posterior distributions are predictions, in order to focus on leaves for which state was predicted with a greater degree of certainty, we now consider the subset of predicted annotations whose 95% credible interval was either entirely below 0.1 or entirely above above 0.9, i.e., low or high chance of having the function respectively. We then compared the list of annotations to those available from the QuickGO API, regardless of the evidence code, which resulted in a list of 295 novel predictions.

Ten of the predictions were for genes in the mouse, a well-studied organism [16]. We searched the literature for evidence of the predicted functions, and uncovered evidence for six of the ten predictions: estradiol dehydrogenase activity for Hsd17b1 [15], involvement in response to iron ion for Slc11a2 [28], involvement of Bok in promoting apoptosis [4], DNA binding activity of Nfib [17], extracellular location for Lipg [1], the adenylate kinase activity of Ak2 [21]. The full list, including the programs used to generate it, is available in this website: https://github.com/USCbiostats/aphylo-simulations.

## 4 Discussion

In this paper, we presented a model for the evolution of gene function that allows rapid inference of that function, along with the associated evolutionary parameters. Such a scheme allows for the hope of *automated* updating of gene function annotations as more experimental information is gathered. We note that our approach results in probabilistic inference of function, thereby capturing uncertainty in a concise and natural way. In simulation studies, using phylogenies from PANTHER, we demonstrate that our approach performs well. Furthermore, our computational implementation of this approach allows for rapid inference across thousands of trees in a short time period.

We believe that basing inference upon an evolutionary model allows us to capture biological properties of gene evolution, and that doing so results in more accurate inference. However, being based on a model also means being “wrong” [2]. We now discuss the ways in which we are wrong, and how we believe that despite this “wrongness” our approach is still useful.

### Branch length

We have treated the probability of a change of function from node-to-node as unrelated to the length of the branch that connects those nodes. This choice follows the prevailing model of gene functional change occurring relatively rapidly after gene duplication, rather than gradually over time. Extension to models in which this is not true is conceptually straightforward, involving a move from discrete to continuous underlying Markov Chains. Parameters *μ*_01_ and *μ*_10_ are then treated as rates rather than probabilities: 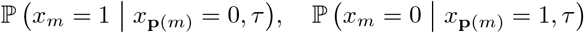, where *τ* is the length of the branch connecting that pair of nodes.

### Multiple gene functions or families

When analyzing multiple gene functions, we have either treated them as if they all had the same parameters values, or were all completely independent. These cases represent two ends of a spectrum, and neither end is likely to be the truth. Even absent any specific knowledge regarding how genes might be related to each other, the sparseness of experimental annotation, and the low frequency of *“no function”* annotations, makes it desirable to take advantage of multiple annotations across functions in order to obtain better parameter inference (see Table 6). If there are *p* gene functions in a particular gene family of interest, a fully saturated model would require 7 × 2*p* parameters (4 sets of mutation rates, 2 sets of misclassification probabilities, and 1 set of root node probabilities). This, clearly, rapidly becomes very challenging, even for “not very large at all” *p*. Thus, more sophisticated approaches will be needed.

**Table 6.** Distribution of trees per number of functions (annotations) in PANTHERDB. At least 50% of the trees used in this paper have 10 or more annotations per tree.

| N Ann. per Tree | N | Cum. count | Cum. prop. |
| --- | --- | --- | --- |
| 1 | 1173 | 1173 | 0.11 |
| 2 | 753 | 1926 | 0.17 |
| 3 | 675 | 2601 | 0.23 |
| 4 | 573 | 3174 | 0.29 |
| 5 | 567 | 3741 | 0.34 |
| 6 | 485 | 4226 | 0.38 |
| 7 | 504 | 4730 | 0.43 |
| 8 | 387 | 5117 | 0.46 |
| 9 | 323 | 5440 | 0.49 |
| 10 or more | 5666 | 11106 | 1.00 |

For now, we believe that the pooled analysis here demonstrates that there is likely to be some utility in developing approaches that can effectively impute annotations for multiple genes or gene functions jointly, particularly when, as is generally the case at present, annotation data is sparse. In future work we intend to explore hierarchical approaches to this problem. For example, it seems reasonable to assume that even if different gene families have different parameter values, the basic evolutionary biology would be similar across all gene families and this could be reflected by modeling the various parameters as random effects in a hierarchical framework. Thus, one would fit the entire ensemble of genes, or gene functions, simultaneously, estimating parameters for their overall means and variances across families. Such an approach would also allow for formal testing for parameter value differences between different families. While such an approach will be challenging to implement, we intend to pursue it in future work.

We also intend to explore approaches in which we use a simple loglinear model to capture the key features. For example, in such an approach we might include *p* parameters for the marginal probabilities and 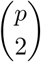 parameters for pairwise associations for the mutation rates and baseline probabilities, for example, while treating the misclassification probabilities as independent. There are many possible extensions of this loglinear framework. For example, it seems likely at after a duplication event, as a function is lost in one branch a new function will be gained in that branch while the other branch remains intact. Such possibilities can be readily incorporated within this framework by modeling gain/loss probabilities jointly.

### Parameters suggest biological interpretations

Our model, while simple, has several advantages and appears to perform well overall. We are able to determine parameters not only for one family at a time, but shared parameters over the set of all 141 protein families that have both positive and negative examples of a given function. The parameters have straightforward interpretations. In agreement with the prevailing model of the importance of gene duplication, the probability of function change (either gain or loss) derived from our model is much larger following gene duplication than following speciation. The high probability of an arbitrary function being present at the root of the tree (*π*) is consistent with the observation that functions are often highly conserved across long evolutionary time spans [20]. Our model also contains some features that may offer additional biological insights. Our sensitivity analysis shows that a small probability of functional loss following speciation is particularly important to prediction accuracy. In other words, functions, once gained, strongly tend to be inherited in the absence of gene duplication. This includes not only molecular functions as generally accepted, but cellular components (the places where proteins are active), and biological processes (larger processes that proteins contribute to) as well.

### Utility of predictions

The predictions from our method may have utility, both in guiding experimental work, or in highlighting areas of conflicting scientific results. High probability predictions are likely to be correct: for the ten such predictions we made for mouse genes, we found experimental evidence for six of them (which were not yet in the GO knowledgebase), and for the remaining four, we found no evidence of either presence or absence of that function. In our leave-one-out predictions of experimentally characterized functions of the EDEM family, we found cases where our predictions were particularly poor, but upon close examination turned out to be indicative of actual conflicts in experimental results from different studies. Deeper analyses of discrepant predictions could be helpful in identifying similar cases.

### Improving the input data using taxon constraints

Another type information that can be leveraged is taxon constraints. Taxon constraints – which we define as a set of biological assumptions that restrict the set of possible values for given nodes on the tree (either by assuming gene function will be present or absent there) – can be used to inform our analysis. Within the context of our model, it is simple to specifying that a clade can or cannot have a particular function, (essentially treating it as if it were fully annotated, without error). Thus, inclusion of such constraints would reduce the uncertainty in the model by effectively decreasing the overall depth of the unannotated parts of the tree (distance from the most recent common ancestor to the tip). More importantly, it would also act to increase the number of available annotations, most likely adding those of type *absent*, which as we show in section 3, are the scarcest ones.

### Use in epidemiologic analyses

We emphasize that the goal of this method is not simply to assign presence or absence of various gene functions to presently unannotated human genes, but to estimate the probabilities *π_gp_* that each gene *g* has each function p. In analyzing epidemiologic studies of gene-disease associations, we anticipate using this annotation information as “prior covariates” in a hierarchical model, in which the first level would be a conventional multiple logistic regression for the log relative risk coefficients *β_g_* for each gene g and the second level would be a regression of these *β_g_* on the vector *π_g_* = (*π_pg_*) of estimated function probabilities. This second level model could take various forms, depending upon whether one thought the functions were informative about both the magnitude and sign of the relative risks or only their magnitudes. In former case, one might adopt a linear regression of the form 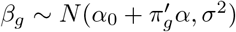. In the latter case, one might instead model their dispersion as *β_g_* ~ *N*(0, λ_*g*_) or *β_g_* ~ Laplace(λ_*g*_) where 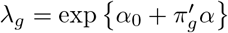, corresponding to an individualized form of ridge or Lasso penalized regression respectively.

In summary, we have presented a parsimonious evolutionary model of gene function. While we intend to further develop this model to reflect additional biological features, we note that in its current form it has the following key features: (a) It is computationally scalable, making it trivial to jointly analyze hundreds of annotated trees in finite time. (b) It yields data-driven results that are aligned with our biological intuition, in particular, supporting the idea that functional changes are most likely to occur during gene-duplication events. (c) Notwithstanding its simplicity, it provides probabilistic predictions with an accuracy level comparable to that of other, more complex, phylo-based models. (d) Perhaps most importantly, it can be used to both support new annotations and to suggest areas in which existing annotations show inconsistencies that may indicate errors or controversies in those experimental annotations.

## 5 Material and Methods

### 5.1 Prediction of Annotations

Let 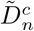 denote an annotated tree with all tree structure and annotations below node *n* removed, the complement of the induced sub-tree of *n*, 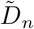. Ultimately we are interested on the conditional probability of the *n*th node having the function of interest given 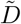, the observed tree and annotations. Let *s* ∈ {0,1}, then we need to compute:

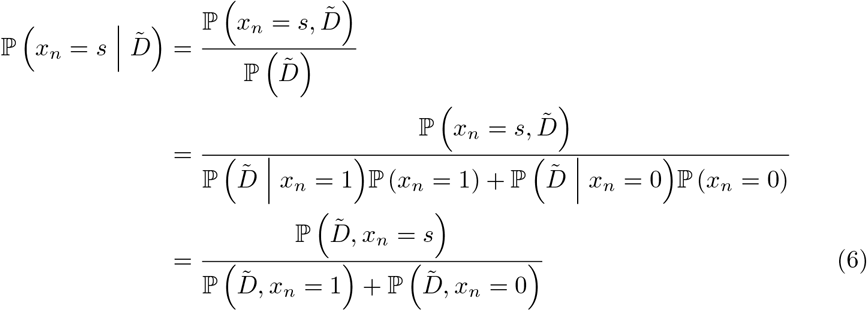

Using conditional independence (which follows from the Markovian property), the joint probability of 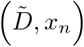 can be decomposed into two pieces, the “pruning probability”, which has already been calculated using the peeling algorithm described in section 2.2.2, and the joint probability of 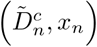:

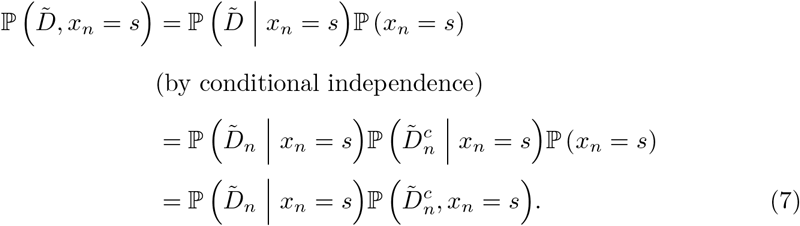

Using the law of total probability, the second term of (7) can be expressed in terms of *n*’s parent state, *x*_**p**(*n*)_, as:

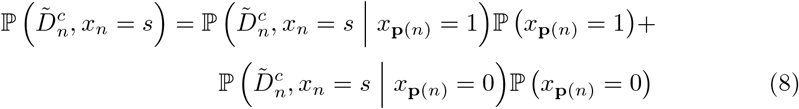

Again, given the state of the parent node, *x*_**p**(*n*)_, *x_n_* and 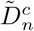 are conditionally independent, and with *s*’ ∈ {0,1}, we have

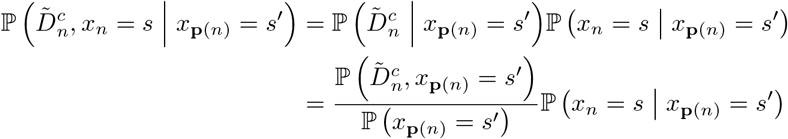

A couple of observations from the previous equation. First, while 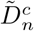 includes node **p** (*n*), it does not include information about its state, *x*_**p**(*n*)_, since only leaf annotations are observed. Second, the equation now includes 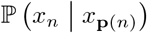, this is, the model’s transition probabilities, (*μ*_01_,*μ*_10_). With the above equation we can write (8) as:

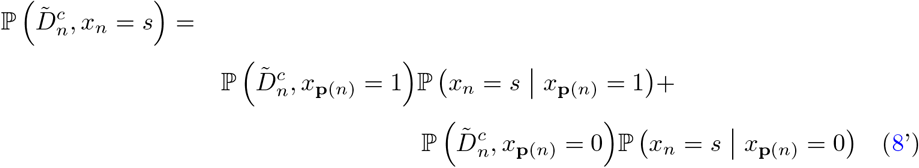

In (8’), the only missing piece is 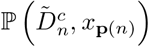, which can be expressed as

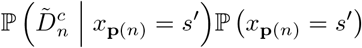

Furthermore, another application of the Markovian property allows us to decompose 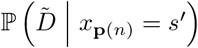 as a product of conditional probabilities. Thus, 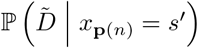 can be expressed as

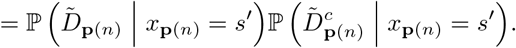

(which, by definition, is)

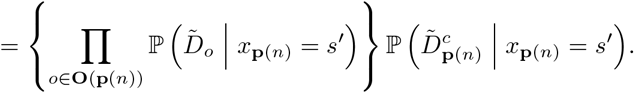

(and, taking the pruning probability of node *n* out of the product operator, gives)

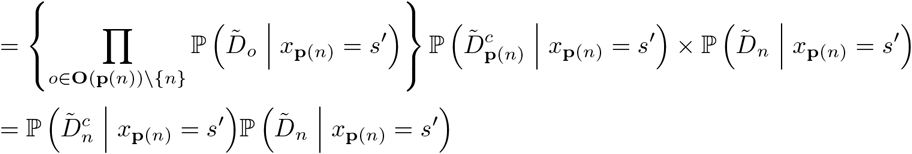

Where the last equality holds by definition of 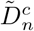. In essence, this shows that we can decompose the conditional probability 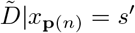 by splitting the tree at either *n* or **p** (*n*). This allows us to calculate 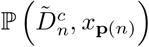:

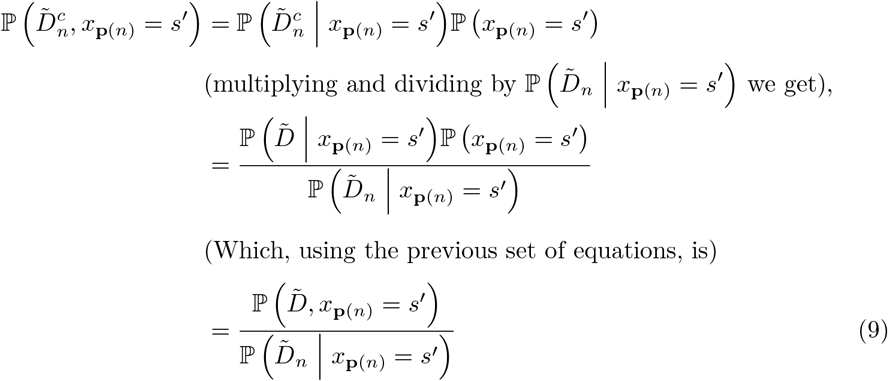

The denominator of (9) can then be rewritten as follows:

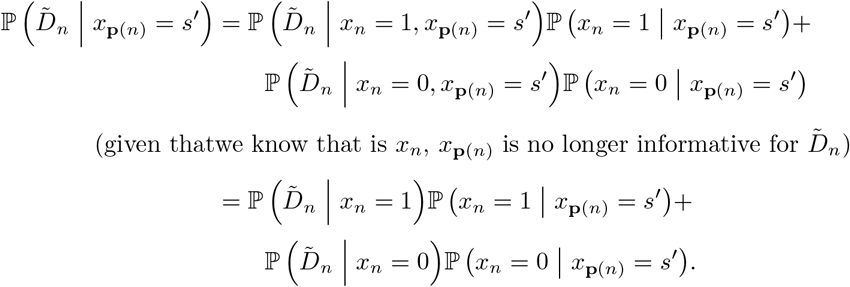

Now, we can substitute this into the denominator of (9) and write:

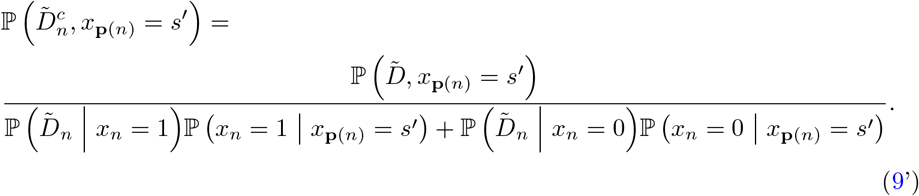

This way, the probability of observing 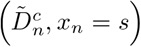, (8’), equals:

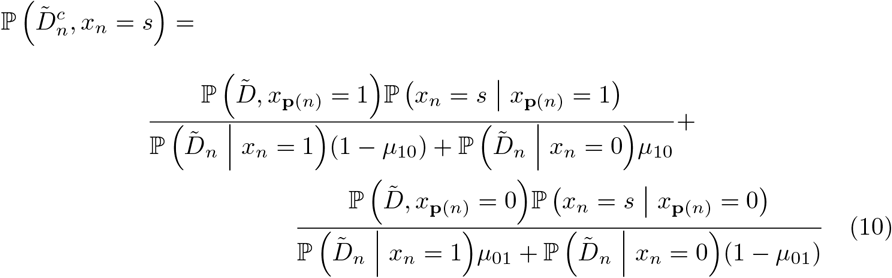

Together with the pruning probabilities calculated during the model fitting process, this equation can be computed using a recursive algorithm, in particular the pre-order traversal [23], in which we iterate through the nodes from root to leaves.

When node *n* is the root node, the posterior probability can be calculated in a straightforward way:

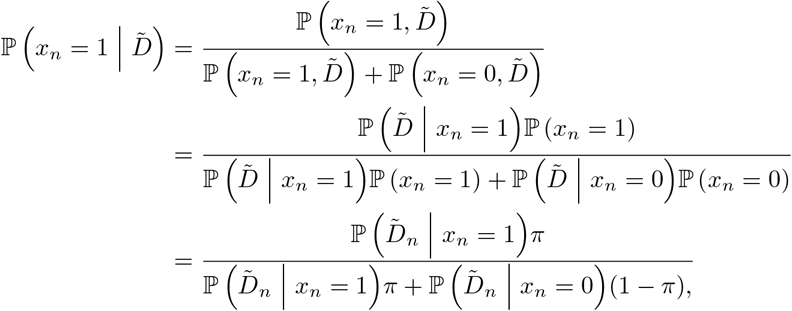

since the terms 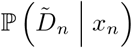 have already been calculated as part of the pruning algorithm.

### 5.2 Monte Carlo Study

We assess model performance by quantifying the quality of our predictions under several scenarios using annotations constructed by simulating the evolutionary process on *real* trees obtained from PANTHER. In particular, we simulate the following scenarios:

1. **Fully annotated**: For each tree in PANTHER we simulated the evolution of gene function, and the annotation of that function at the tree tips, using the model described in this paper. For each tree we drew a different set of model parameters from the following Beta distribution: We then used the simulated annotations to estimate the model parameters and gene function states. For this case we exclude mislabeling, i.e. all leaves are correctly annotated.

2. **Partially annotated**: Here, we took the set of simulations produced in scenario 1 above, but now estimated the model using a partially annotated tree. Specifically, we randomly dropped a proportion of leaf annotations *m* ~ Uniform(0.1,0.9). Once again, we assumed no mislabeling. (So, *ψ***_01_** = 0, *ψ***_10_** = 0).
3. **Partially annotated with mislabeling** Finally, we take the data from scenario 2 but allow for the possibility of mislabeling in the annotations. Specifically, for each tree we draw values for *ψ*_01_ and *ψ*_10_ from a Beta(2, 38) distribution.

| Parameter | $\alpha$ | $\beta$ | Mean<br>$\alpha/(\alpha + \beta)$ |
| --- | --- | --- | --- |
| $\mu_{01d}$ | 38 | 2 | 0.95 |
| $\mu_{10d}$ | 10 | 10 | 0.50 |
| $\mu_{01s}, \mu_{10s}$ | 2 | 38 | 0.05 |
| $\pi$ | 2 | 38 | 0.05 |

In order to assess the effect of the prior distribution, in each scenario we performed estimation twice using two different priors: a well-specified prior, i.e., the one used during the data-generating-process, and a biased prior in which the *α* shape was twice of that of the data-generating-process.

#### 5.2.1 Accuracy

Figure 6 shows the distribution of AUCs and Mean Absolute Errors [MAEs] for the third scenario by prior used and proportion of missing labels.

**Fig 6.**
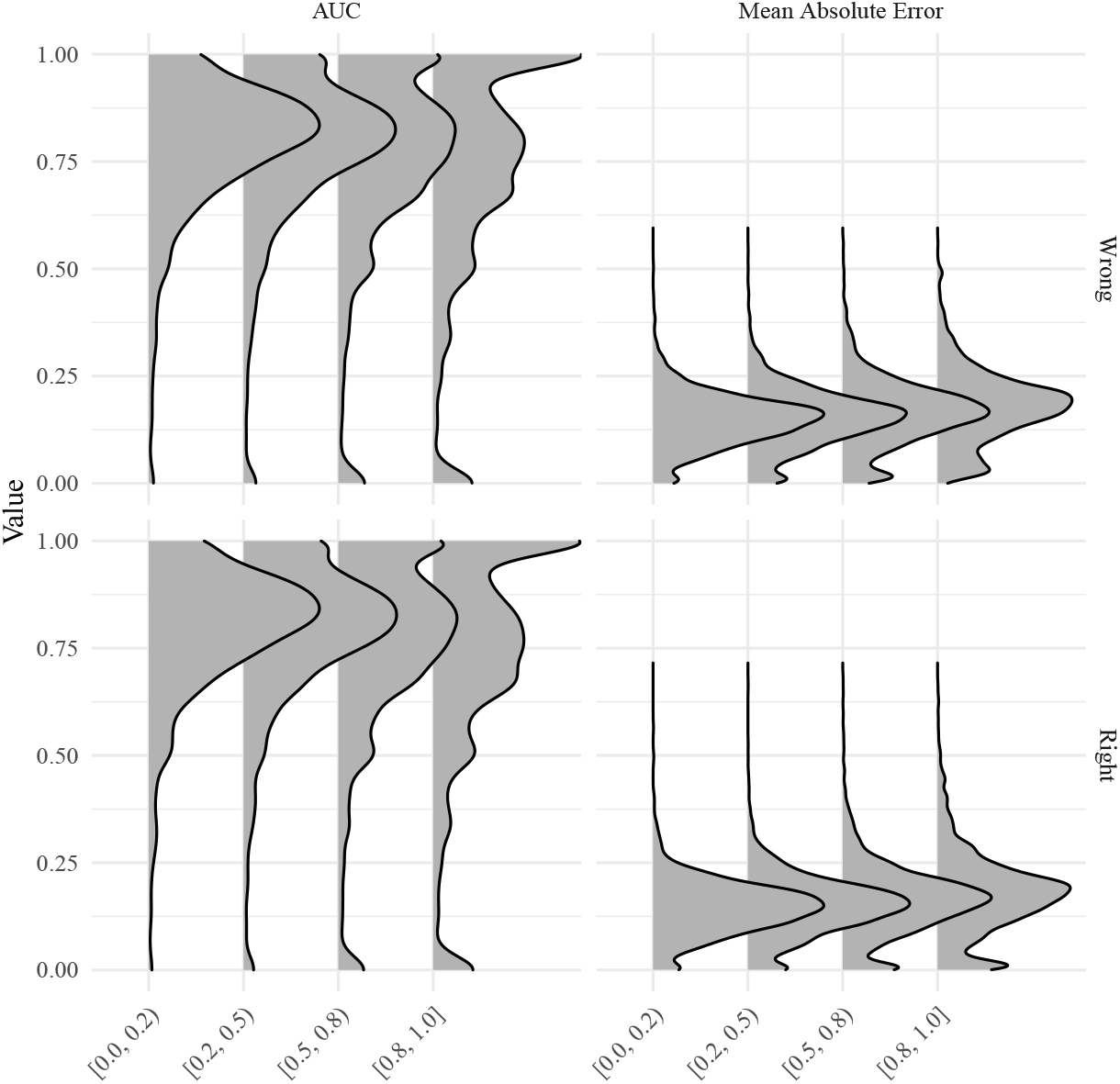
Distribution of AUCs and MAEs for the scenario with partially annotated trees and mislabeling. The x-axis shows the proportion of missing annotations, while the y-axis shows the score (AUC or MAE).

Overall, the predictions are good with a relatively low MAE/high AUC. Furthermore, as seen in Figure 6, both AUC and MAE worsen off as the data becomes more sparse (fewer annotated leaves).

#### 5.2.2 Bias

We now consider bias. Figure 7 shows the distribution of the empirical bias, defined as the population parameter minus the parameter estimate, for the first scenario (fully annotated tree). Since the tree is fully annotated and there is no mislabeling, the plot only shows the parameters for functional gain, loss and the root node probability. Of the three parameters, *π* is the one which the model has the hardest time to recover, what’s more, it generally seems to be over-estimated. On the other hand, *μ*_01_,*μ***10** estimates do significantly better than those for *π*, and moreover, in sufficiently large trees the model with the correct prior is able to recover the true parameter value.

**Fig 7.**
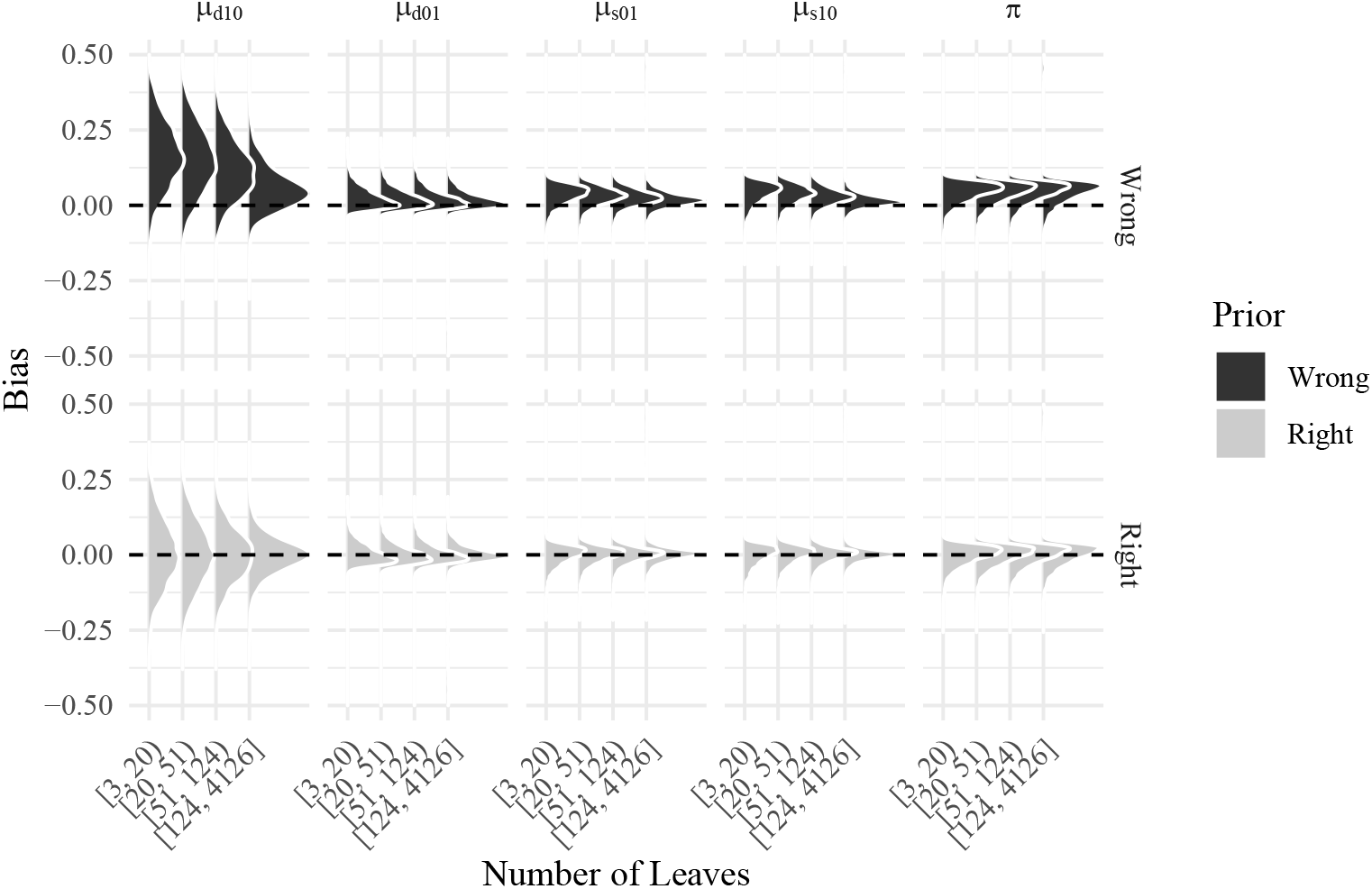
Empirical bias distribution for the Fully annotated scenario by type of prior, parameter, and number of leaves.

Figure 8 shows the empirical distribution of the parameter estimates in the third simulation scheme: a partially annotated tree with mislabelling. As the proportion of missing annotations increases, the model tends to, as expected, do worse.

**Fig 8.**
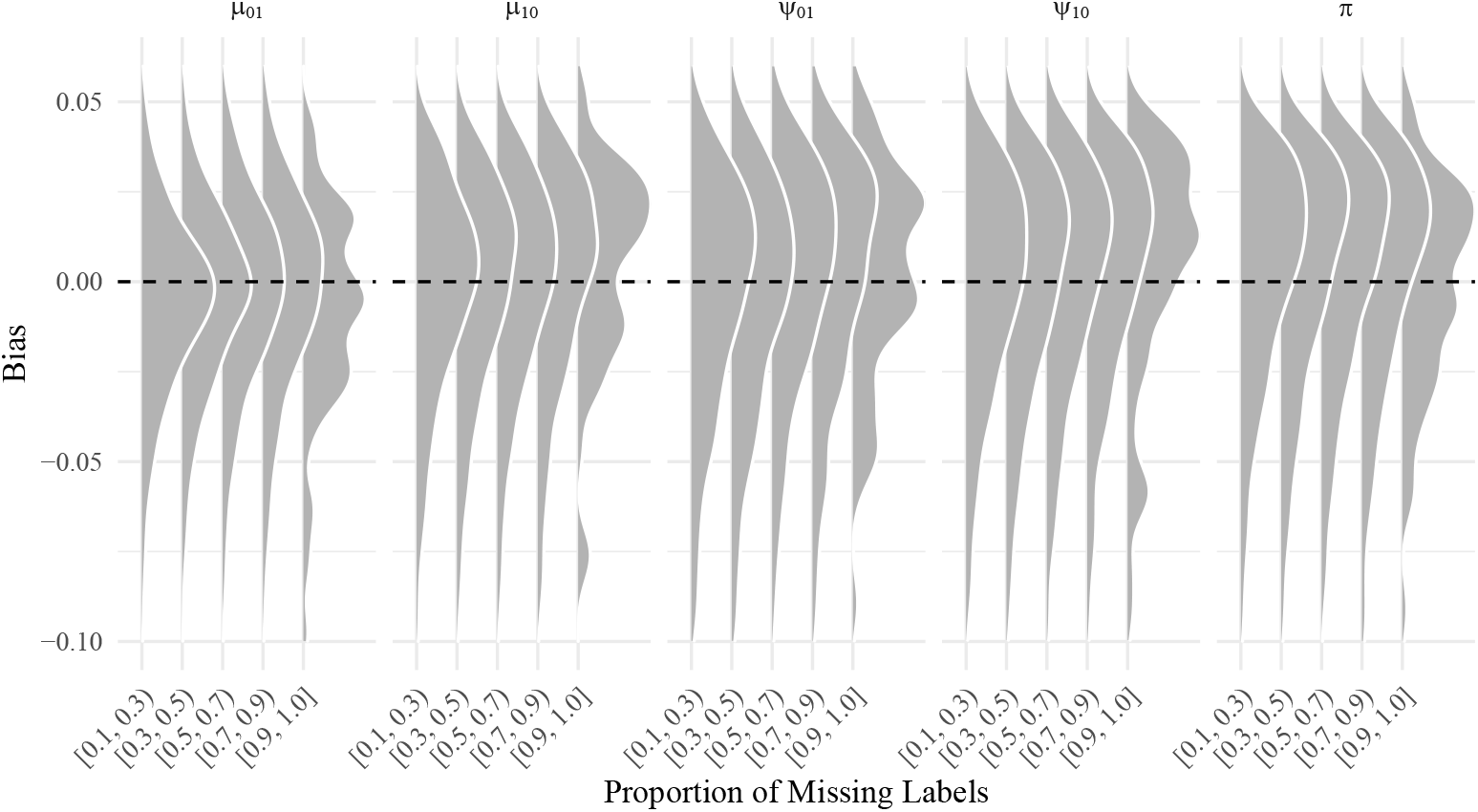
Empirical bias distribution for the partially annotated scenario by parameter and proportion of missing labels.

### 5.3 Limiting probabilities

In the case that the model includes a single set of gain and loss probabilities, (i.e. no difference according to type of node), in the limit as the number of branches between the root and the node goes to infinity, the probability that a given node has a function can be calculated as

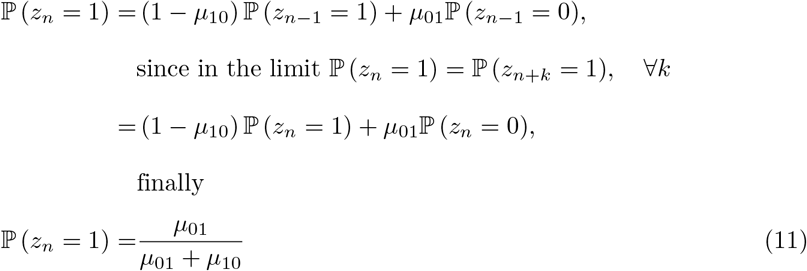

However, when the gain and loss probabilities differ by type of node, the unconditional probability of observing having function is computed as follows:

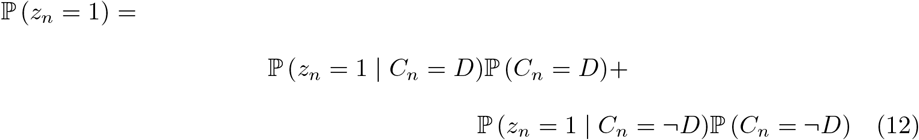

Where 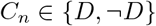 denotes the type of event (duplication or not duplication, respectively). We need to calculate 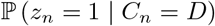 and 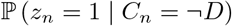. WLOG let’s start by the first:

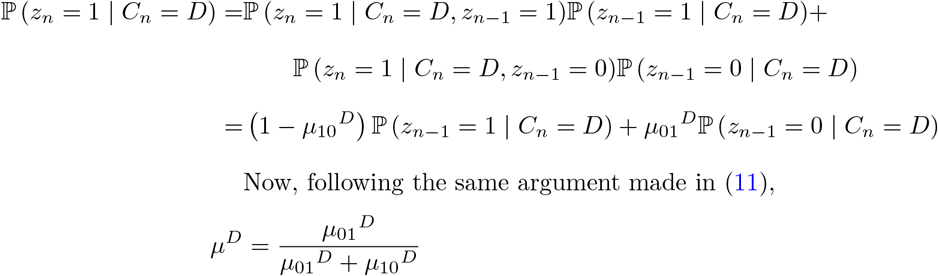

Where 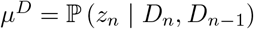. Likewise, 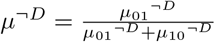. Observe that the parameter is only indexed by the class of the *n*-th leaf as the class of its parent is not relevant for this calculation.

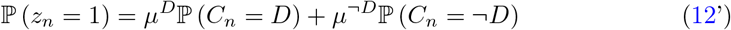

Where the probability of a node having a given class can be approximated by the observed proportion of that class in the given tree.

## Acknowledgements

Supported by National Cancer Institute Grant #1P01CA196596. Computation for the work described in this paper was supported by the University of Southern California’s Center for High-Performance Computing (https://hpcc.usc.edu).

## References

[1] Debapriya Basu et al. “Measurement of the phospholipase activity of endothelial lipase in mouse plasma”. In: Journal of Lipid Research 54.1 (Jan. 2013), pp. 282–289. ISSN: 0022-2275. DOI: 10.1194/jlr.D031112. URL: http://www.jlr.org/lookup/doi/10.1194/jlr.D031112.

[2] George E. P. Box. “Science and Statistics”. In: Journal of the American Statistical Association 71.356 (Dec. 1976), pp. 791–799. ISSN: 0162-1459. DOI: 10.1080/01621459.1976.10480949. URL: http://www.tandfonline.com/doi/abs/10.1080/01621459.1976.10480949.

[3] S Brooks et al. Handbook of Markov Chain Monte Carlo. Chapman & Hall/CRC Handbooks of Modern Statistical Methods. CRC Press, 2011. ISBN: 9781420079425. URL: https://books.google.com/books?id=qfRsAIKZ4rIC.

[4] Marcos A. Carpio et al. “BCL-2 family member BOK promotes apoptosis in response to endoplasmic reticulum stress”. In: Proceedings of the National Academy of Sciences 112.23 (June 2015), pp. 7201–7206. ISSN: 0027-8424. DOI: 10.1073/pnas.1421063112. URL: http://www.pnas.org/lookup/doi/10.1073/pnas.1421063112.

[5] Davide Chicco. “Ten quick tips for machine learning in computational biology”. In: BioData Mining 10.1 (Dec. 2017), p. 35. ISSN: 1756-0381. DOI: 10.1186/s13040-017-0155-3. URL: https://biodatamining.biomedcentral.com/articles/10.1186/s13040-017-0155-3.

[6] Jonathan A. Eisen. “Phylogenomics: Improving Functional Predictions for Uncharacterized Genes by Evolutionary Analysis”. In: Genome Research 8.3 (1998), pp. 163–167. DOI: 10.1101/gr.8.3.163. eprint: http://genome.cshlp.org/content/8/3/163.full.pdf+html: http://genome.cshlp.org/content/8/3/163.short.

[7] Barbara E Engelhardt et al. “Genome-scale phylogenetic function annotation of large and diverse protein families”. In: Genome research 21.11 (2011), pp. 1969–1980. DOI: 10.1101/gr.104687.109.

[8] Barbara E Engelhardt et al. “Protein Molecular Function Prediction by Bayesian Phylogenomics”. In: PLOS Computational Biology 1.5 (2005). DOI: 10.1371/journal.pcbi.0010045. URL: https://doi.org/10.1371/journal.pcbi.0010045.

[9] Tom Fawcett. “An introduction to ROC analysis”. In: Pattern Recognition Letters 27.8 (June 2006), pp. 861–874. ISSN: 01678655. DOI: 10.1016/j.patrec.2005.10.010. URL: https://linkinghub.elsevier.com/retrieve/pii/S016786550500303X.

[10] Joseph Felsenstein. “Evolutionary trees from DNA sequences: A maximum likelihood approach”. In: Journal of Molecular Evolution 17.6 (Nov. 1981), pp. 368–376. ISSN: 1432-1432. DOI: 10.1007/BF01734359. URL: https://doi.org/10.1007/BF01734359.

[11] C. Ferri, J. Hernandez-Orallo, and R. Modroiu. “An experimental comparison of performance measures for classification”. In: Pattern Recognition Letters 30.1 (Jan. 2009), pp. 27–38. ISSN: 01678655. DOI: 10.1016/j.patrec.2008.08.010. URL: https://linkinghub.elsevier.com/retrieve/pii/S0167865508002687.

[12] Olivier Gascuel and Mike Steel. “Predicting the Ancestral Character Changes in a Tree is Typically Easier than Predicting the Root State”. In: Systematic Biology 63.3 (May 2014), pp. 421–435. ISSN: 1076-836X. DOI: 10.1093/sysbio/syu010. URL: https://academic.oup.com/sysbio/article-lookup/doi/10.1093/sysbio/syu010.

[13] Pascale Gaudet et al. “Phylogenetic-based propagation of functional annotations within the Gene Ontology consortium”. In: Briefings in Bioinformatics 12.5 (2011), pp. 449–462. DOI: 10.1093/bib/bbr042. eprint: //oup/backfile/content_public/journal/bib/12/5/10.1093/bib/bbr042/2/bbr042.pdf. URL: +%20http://dx.doi.org/10.1093/bib/bbr042.

[14] Heikki Haario, Eero Saksman, and Johanna Tamminen. “An adaptive Metropolis algorithm”. In: Bernoulli 7.2 (2001), pp. 223–242. URL: https://projecteuclid.org/euclid.bj/1080222083.

[15] Janne Hakkarainen et al. “Hydroxysteroid (17β) dehydrogenase 1 expressed by Sertoli cells contributes to steroid synthesis and is required for male fertility”. In: The FASEB Journal 32.6 (June 2018), pp. 3229–3241. ISSN: 0892-6638. DOI: 10.1096/fj.201700921R. URL: https://onlinelibrary.wiley.com/doi/abs/10.1096/fj.201700921R.

[16] Douglas G. Howe et al. “Model organism data evolving in support of translational medicine”. In: Lab Animal 47.10 (Oct. 2018), pp. 277–289. ISSN: 0093-7355. DOI: 10.1038/s41684-018-0150-4. URL: http://www.nature.com/articles/s41684-018-0150-4.

[17] Yu-Chih Hsu et al. “Mesenchymal Nuclear factor I B regulates cell proliferation and epithelial differentiation during lung maturation”. In: Developmental Biology 354.2 (June 2011), pp. 242–252. ISSN: 00121606. DOI: 10.1016/j.ydbio.2011.04.002. URL: https://linkinghub.elsevier.com/retrieve/pii/S0012160611002120.

[18] Sohta A Ishikawa et al. “A Fast Likelihood Method to Reconstruct and Visualize Ancestral Scenarios”. In: Molecular Biology and Evolution 36.9 (Sept. 2019). Ed. by Tal Pupko, pp. 2069–2085. ISSN: 0737-4038. DOI: 10.1093/molbev/msz131. URL: https://academic.oup.com/mbe/article/36/9/2069/5498561.

[19] Yuxiang Jiang et al. “An expanded evaluation of protein function prediction methods shows an improvement in accuracy”. In: Genome Biology 17.1 (Sept. 2016), p. 184. ISSN: 1474-760X. DOI: 10.1186/s13059-016-1037-6. URL: https://doi.org/10.1186/s13059-016-1037-6.

[20] A. H. Kachroo et al. “Systematic humanization of yeast genes reveals conserved functions and genetic modularity”. In: Science 348.6237 (May 2015), pp. 921–925. ISSN: 0036-8075. DOI: 10.1126/science.aaa0769. URL: https://www.sciencemag.org/lookup/doi/10.1126/science.aaa0769.

[21] Aleksandr Klepinin et al. “Simple oxygraphic analysis for the presence of adenylate kinase 1 and 2 in normal and tumor cells”. In: Journal of Bioenergetics and Biomembranes 48.5 (Oct. 2016), pp. 531–548. ISSN: 0145-479X. DOI: 10.1007/s10863-016-9687-3. URL: http://link.springer.com/10.1007/s10863-016-9687-3.

[22] Huaiyu Mi et al. “PANTHER version 11: expanded annotation data from Gene Ontology and Reactome pathways, and data analysis tool enhancements”. In: Nucleic Acids Research 45.D1 (2017), pp. D183–D189. DOI: 10.1093/nar/gkw1138. eprint: /oup/backfile/content_public/journal/nar/45/d1/10.1093_nar_gkw1138/3/gkw1138.pdf. URL: +%20http://dx.doi.org/10.1093/nar/gkw1138.

[23] Joseph M. Morris. “Traversing binary trees simply and cheaply”. In: Information Processing Letters 9.5 (Dec. 1979), pp. 197–200. ISSN: 00200190. DOI: 10.1016/0020-0190(79)90068-1. URL: http://linkinghub.elsevier.com/retrieve/pii/0020019079900681.

[24] Satoshi Ninagawa et al. “EDEM2 initiates mammalian glycoprotein ERAD by catalyzing the first mannose trimming step”. In: Journal of Cell Biology 206.3 (Aug. 2014), pp. 347–356. ISSN: 1540-8140. DOI: 10.1083/jcb.201404075. URL: https://rupress.org/jcb/article/206/3/347/37772/EDEM2-initiates-mammalian-glycoprotein-ERAD-by.

[25] E. Paradis and K. Schliep. “ape 5.0: an environment for modern phylogenetics and evolutionary analyses in R”. In: Bioinformatics 35 (2018), pp. 526–528.

[26] R Core Team. R: A Language and Environment for Statistical Computing. Vienna, Austria, 2012. URL: http://www.r-project.org/.

[27] Predrag Radivojac et al. “A large-scale evaluation of computational protein function prediction”. In: Nature Methods 10 (Jan. 2013), p. 221. URL: http://dx.doi.org/10.1038/nmeth.2340%20http://10.0.4.14/nmeth.2340%20https://www.nature.com/articles/nmeth.2340%7B%5C#%7Dsupplementary-information.

[28] Ali Shawki et al. “Intestinal brush-border Na + /H + exchanger-3 drives H + -coupled iron absorption in the mouse”. In: American Journal of Physiology-Gastrointestinal and Liver Physiology 311.3 (Sept. 2016), G423–G430. ISSN: 0193-1857. DOI: 10.1152/ajpgi.00167.2016. URL: https://www.physiology.org/doi/10.1152/ajpgi.00167.2016.

[29] The Gene Ontology Consortium. “Expansion of the Gene Ontology knowledgebase and resources”. In: Nucleic Acids Research 45.D1 (2017), pp. D331–D338. DOI: 10.1093/nar/gkw1108. eprint: //oup/backfile/content_public/journal/nar/45/d1/10.1093_nar_gkw1108/3/gkw1108.pdf. URL: +%20http://dx.doi.org/10.1093/nar/gkw1108.

[30] George Vega Yon and Paul Marjoram. “fmcmc: A friendly MCMC framework”. In: Journal of Open S’ource Software 4.39 (July 2019), p. 1427. ISSN: 2475-9066. DOI: 10.21105/joss.01427. URL: http://joss.theoj.org/papers/10.21105/joss.01427.

[31] Hadley Wickham. ggplot2: Elegant Graphics for Data Analysis. Springer-Verlag New York, 2016. ISBN: 978-3-319-24277-4. URL: https://ggplot2.tidyverse.org.

[32] Ziheng Yang and Carlos E. Rodríguez. “Searching for efficient Markov chain Monte Carlo proposal kernels”. In: Proceedings of the National Academy of Sciences 110.48 (2013), pp. 19307–19312. DOI: 10.1073/pnas.1311790110. eprint: http://www.pnas.org/content/110/48/19307.full.pdf. URL: http://www.pnas.org/content/110/48/19307.abstract.

